# Artifact Formation in Single Molecule Localization Microscopy

**DOI:** 10.1101/700955

**Authors:** Jochen M. Reichel, Thomas Vomhof, Jens Michaelis

## Abstract

We investigate the influence of different accuracy-detection rate trade-offs on image reconstruction in single molecule localization microscopy. Our main focus is the investigation of image artifacts experienced when using low localization accuracy, especially in the presence of sample drift and inhomogeneous background. In this context we present a newly developed SMLM software termed FIRESTORM which is optimized for high accuracy reconstruction. For our analysis we used *in silico* SMLM data and compared the reconstructed images to the ground truth data. We observe two discriminable reconstruction populations of which only one shows the desired localization behavior.

## Introduction

Single Molecule Localization Microscopy (SMLM) is an umbrella term for a multitude of far-field super-resolution techniques whose common features are the optical isolation of single emitter point-spread-functions (PSFs) and the subsequent determination of the respective emitter position. The precision of single emitter localization is not limited by classical microscopy resolution limits but primarily by the number of detected photons [1]. Hence, it is possible to reconstruct a super-resolved image from an adequate number of localizations. PALM [2], FPALM [3], STORM [4], dSTORM [5] and PAINT [6] are only some of the techniques considered to fall under the term SMLM [7] [8] [9] [10] [11]. With a spatial localization precision of typically few tens of nanometers within biological samples, SMLM has become a vital part of the life sciences in the last decade [12] [13] [14] [15] [16] [17].

The localization of emitter positions from spatial intensity distributions is the corner stone of SMLM. Typically the distribution is compared to a model function either via least-square fitting or maximum likelihood estimation, though other approaches are used as well [18] [19] [20] [21]. Here, especially machine learning approaches are becoming increasingly more widespread [22] [23] [24]. Every localization is characterized by its individual localization precision, usually expressed in terms of the localization standard deviation *σ* or the localization full-width-at-half-maximum (FWHM). It is self-evident that the average localization precision of a SMLM image can be improved by only reconstructing localizations that are likely to have a high precision e.g. by choosing localizations based on a high number of photons. Setting such a threshold improves the precision at the expense of the labelling recall which is a common trade-off in SMLM [25] [26]. However, the structure under investigation i.e. its highest spatial frequency sets a lower limit for the necessary number of localizations used in its reconstruction. An oversampling factor of at least fivefold the number of localizations necessary to fulfill the Nyquist criterion was suggested [27]. Undersampling of any structure will deteriorate the image resolution regardless of the average localization precision [28] [27]. Thus, the determination of the appropriate recall-precision trade-off is crucial for the resolution of the reconstruction.

Inappropriate fitting of PSFs and defining overly relaxed thresholds for localization acceptance give rise to image reconstructions that do not properly reflect the structure under investigation. Ultimately, the formation of these reconstruction artifacts might lead to misinterpretation of the acquired data. Thus, we aim to investigate the nature of those low accuracy artifacts. The influence of thermal drift and background correction is discussed as well. Localization microscopy artifacts and their formation is a topic already extensively covered in the SMLM literature [29] [30] [31] [32]. We also present a new analysis software - FIRESTORM - that is optimized for high accuracy reconstructions of SMLM data.

## Materials and Methods

### Data Simulation

#### Simulation without drift

In order to analyze accuracy and precision of SMLM, simulated data based on a ground truth image is required. The chosen ground truth image of 384 x 384 px^2^ with a simulated pixel size of 10 nm is shown in Fig 1A. The simulated structures are a large cross (upper left corner), a series of parallel, single pixel lines with a spacing of 20 nm, 30 nm, 40 nm, 50 nm, 100 nm and 200 nm (from left to right), a second cross whose lines meet under an acute angle, two circles with a diameter of 100 nm and a line width of 10 nm and 20 nm respectively and four lines with a labeling density of 25%, 50%, 75% and 100%. For visualization purposes, the labeling density is encoded as the gray value of the described structures. These structures were not chosen to mimic biological structures but to facilitate a basic analysis of artifact formation in SMLM.

**Fig 1.**
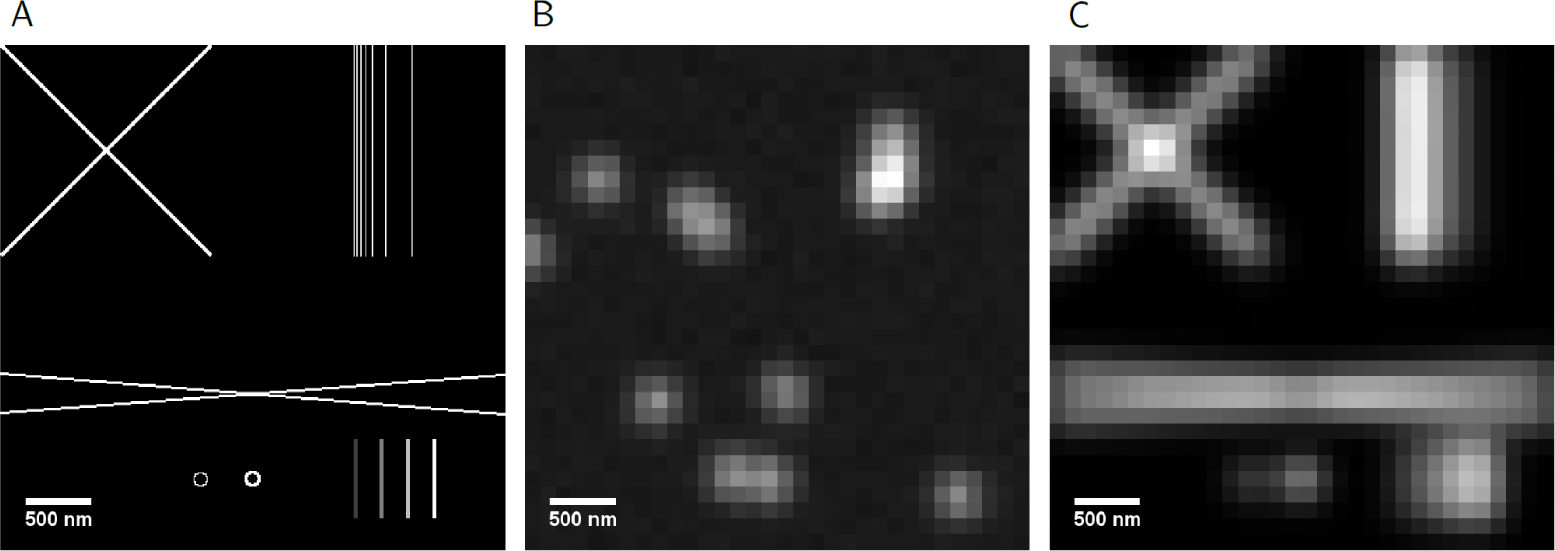
Simulated SMLM data. A: Ground truth image with a pixel size of 10 nm. B: Single frame of the simulated data set displaying individual blinking events. C: Average over the entire image set of 10000 frames; equivalent to the wide field image.

Based on the ground truth image a series of 10000 frames is generated within a custom written MATLAB program (Supplementary Data). The FWHM of the simulated PSF is 300 nm with an average number of 10 blinking events per frame. Over the entire image set 99708 ground truth blinking events are simulated. A single event is modeled to contain an average number of 3000 detected photons, while the average background level is defined as 50 photons. Signal and background photon numbers are both distributed according to Poisson statistics. The detection is modeled to simulate an EMCCD camera. Each low resolution camera frame has a size of 32 x 32 px^2^ with a pixel size of 120 nm. The EM-gain is set to 100, the sensitivity is 11.2 electrons per count, the base level is 400 counts and the maximum camera count value is 16383.

#### Simulation with drift

Furthermore, we want to simulate lateral drift of the specimen relative to the detection system which is a common problem encountered in microscopy techniques with a high spatial resolution and requiring a long acquisition time. Thus, we simulated a second dataset by overlaying an high resolution image series of blinking events with a non-linear movement in *x* and *y* with a total displacement of about 200 nm over the entire image series before converting it into the respective low resolution image series. Over the entire stack 87872 ground truth blinking events take place. Also, a simulated fiducial marker is added that does not blink but emits continuously with an average photon number of 8000 per frame.

### The FIRESTORM analysis tool

FIRESTORM (**F**luorescent spot **i**dentification and **re**construction of **STORM** data) is a localization microscopy data analysis and reconstruction tool. The software is implemented in MATLAB using parallel computation to increase analysis speed. After optional background correction by a running median temporal filter [33] regions of interest around intensity peaks are determined if the peaks meet the minimal SNR. Within those regions of interest a two-dimensional Gaussian function is fitted to the intensity distribution. The fit is accepted if the user defined threshold are met (that is maximum FWHM, symmetry constant, SNR, minimum number of photons). Fluorescent on times spanning consecutive frames can be combined to a single blinking event with a higher number of photons. Besides the Gaussian fitting there is also the option to use a center of mass modality which allows for almost real time reconstruction but yields inferior results in terms of localization precision.

The localization list is analyzed to determine the distributions of SNR and PSF width as well as the photon statistics. Based on this analysis the localization list can be filtered prior to the reconstruction to yield optimal trade-off between recall and localization precision. The intensity values of the reconstructed image can be chosen to be based on the number of photons (NOP) or the number of localizations (NOL). Drift can be corrected by FIRESTORM either via Redundant Cross Correlation [34] (RCC) or by using fiducial marker positions [35]. The software has additional functionality such as multicolor analysis, not covered here.

### Image Reconstruction

Today, a plethora of SMLM analysis, localization and reconstruction tools are available. For a systematic comparison see the publications of Sage *et al.* [25] [26]. Within our discussion we will focus on QuickPALM [36], the first openly available SMLM software, ThunderSTORM [37], a tool that is very widespread in the SMLM community [38] [39] [40] and FIRESTORM. While QuickPALM uses a center-of-mass approach, ThunderSTORM offers a multitude of fitting and (post-) processing modalities. We used these two programs and FIRESTORM to reconstruct the same simulated SMLM data set. To simplify the comparison of reconstruction artifact to FIRESTORM we used ThunderSTORM in the integrated 2D Gaussian fit modality. For all programs we used identical parameters or default parameters respectively when there was no appropriate equivalent. The upper threshold for the PSF FWHM was set to 360 nm (3 px) which equals a standard deviation of the fitted Gaussian distribution of 153 nm. PSFs with larger FWHM are considered to originate from multiple emitter overlap and are thus excluded. The reconstructions were rendered in the respective tool provided by the software. Pixel intensity is based on the number of localizations but a custom colourmap (black - red - yellow - white) is used that already shows single localizations in red to enhance the visibility of artifacts. In a post-processing step a Gaussian blur with a FWHM of 10 nm is applied and the contrast was adjusted in an identical manner for all reconstructions. The quantitative analysis was performed on the original, unprocessed data.

### Reconstruction evaluation

The quantification of reconstruction quality is based on a comparison between reconstruction and ground truth data as described by Sage *et al*. Thus, we can compare the number of simulated events *S*, the number of localizations *L* and the number of matched pairs *M*. We consider a localization to be a match if it is situated closer than 1.5 px=180 nm (half the upper threshold for the FWHM) to a simulated event. Only the closest true event is considered a match if there is more than one simulated event within that radius around a localization. By comparing the matched pairs to the total number of localizations, including those that are not associated to a simulated event, one can determine the correctness of a reconstruction.

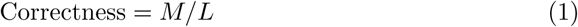

The recall or sensitivity determines what fraction of the simulated events was properly reconstructed.

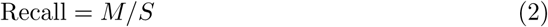

The Jaccard index is a metric that combines correctness and recall:

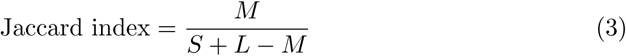

The metrices correctness, recall and the Jaccard index quantify how well an algorithm performs in terms of event identification. To quantify the performance in the next important step, i.e. localization, one typically uses the root-mean-square displacement (RMSD). It compares *N* simulated event positions (*x*^Ref^, *y*^Ref^) to its matched fitted position (*x*^Test^, *y*^Test^).

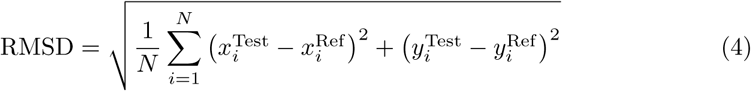

As pointed out by Sage *et al.* it is common practice in the terminology of the SMLM community to use RMSD synonymously with ‘accuracy’ which is misleading in the context of unbiased estimators. Here, we will use the term ‘high accuracy’ to indicate a measurement of high precision **and** high trueness (absence of bias).

## Results and Discussion

### Software comparison

We designed several test structures and simulated realistic SMLM data using these test structures (Materials and Methods). QuickPALM, ThunderSTORM and FIRESTORM all recover the basic features of the ground truth image. As shown in table 1 the programs perform almost identically in terms of correctness. For ThunderSTORM and FIRESTORM the values are at unity, indicating that a reconstructed event always is associated with a real event. Also for QuickPALM only 0.4% of localizations L could not be matched properly to a simulated event. However, there are considerable differences in terms of recall. While QuickPALM (29.8%) and FIRESTORM (30.8%) reconstruct a similar number of events, ThunderSTORM finds 41.9% of the simulated events. Since there is hardly any difference in terms of correctness between the programs, the Jaccard index follows the recall very closely.

**Table 1.** Summary of the quantitative reconstruction comparison between QuickPALM, ThunderSTORM and FIRESTORM for the simulation without drift. Metrics of interest are the number of localizations *L*, the number of matches *M*, the correctness, the recall, the Jaccard index and the root-mean-square displacement. The number of simulated events *S* is 99708.

| | $L$ | $M$ | correctness<br>/ % | recall<br>/ % | Jaccard<br>index / % | RMSD<br>/ nm |
| --- | --- | --- | --- | --- | --- | --- |
| QuickPALM | 29824 | 29690 | 99.6 | 29.8 | 29.7 | 55.5 |
| ThunderSTORM | 41757 | 41757 | 100.0 | 41.9 | 41.9 | 33.2 |
| FIRESTORM | 30750 | 30750 | 100.0 | 30.8 | 30.8 | 21.9 |

With 55.5 nm QuickPALM has the highest RMSD value. That can be explained in large part by the fact that it uses a simple center of mass fitting routine. ThunderSTORM yields a RMSD value of 33.2 nm and FIRESTORM yields 21.9 nm. The better precision obtained by FIRESTORM goes hand in hand with the worse recall, a trade-off commonly observed when comparing SMLM programs. The most straightforward way to tune this trade off is by setting quality thresholds a PSF has to fulfill, e.g. minimum number of photons, minimum SNR, max FWHM or geometrical restrictions like the maximum acceptable ellipticity. By excluding events that do not meet those criteria, it is possible to improve the precision at the expense of the recall. However, choosing overly restrictive thresholds results in reconstructions in which features of interest are undersampled which defeats the purpose of SMLM. The correct parameter choice is thus a crucial step in SMLM data analysis.

### Artifact formation

While all three analysis softwares were able to reconstruct the different designed patterns, some of the images (Fig 2 i) showed additional features not visible in the ground truth image (Fig 1 A), hence they are reconstruction artifacts. Observed artifacts are the blurring of structures at crossing points (best visible at the large cross in the upper, left corner), the formation of ghost lines between the ground truth lines. Also, intensity modulations not present in the homogeneously labelled ground truth image are observed (best visible in the QuickPALM reconstruction of the steep angle cross). In order to quantify the artifact formation we plot the intensity profile of the upper right structure along the *x*-axis with the true position of the seven vertical lines indicated in red. A general loss of intensity modulation depth in regions of fine spatial structures alongside an impaired trueness of the reconstructed line position is observed, especially in the QuickPALM reconstruction.

**Fig 2.**
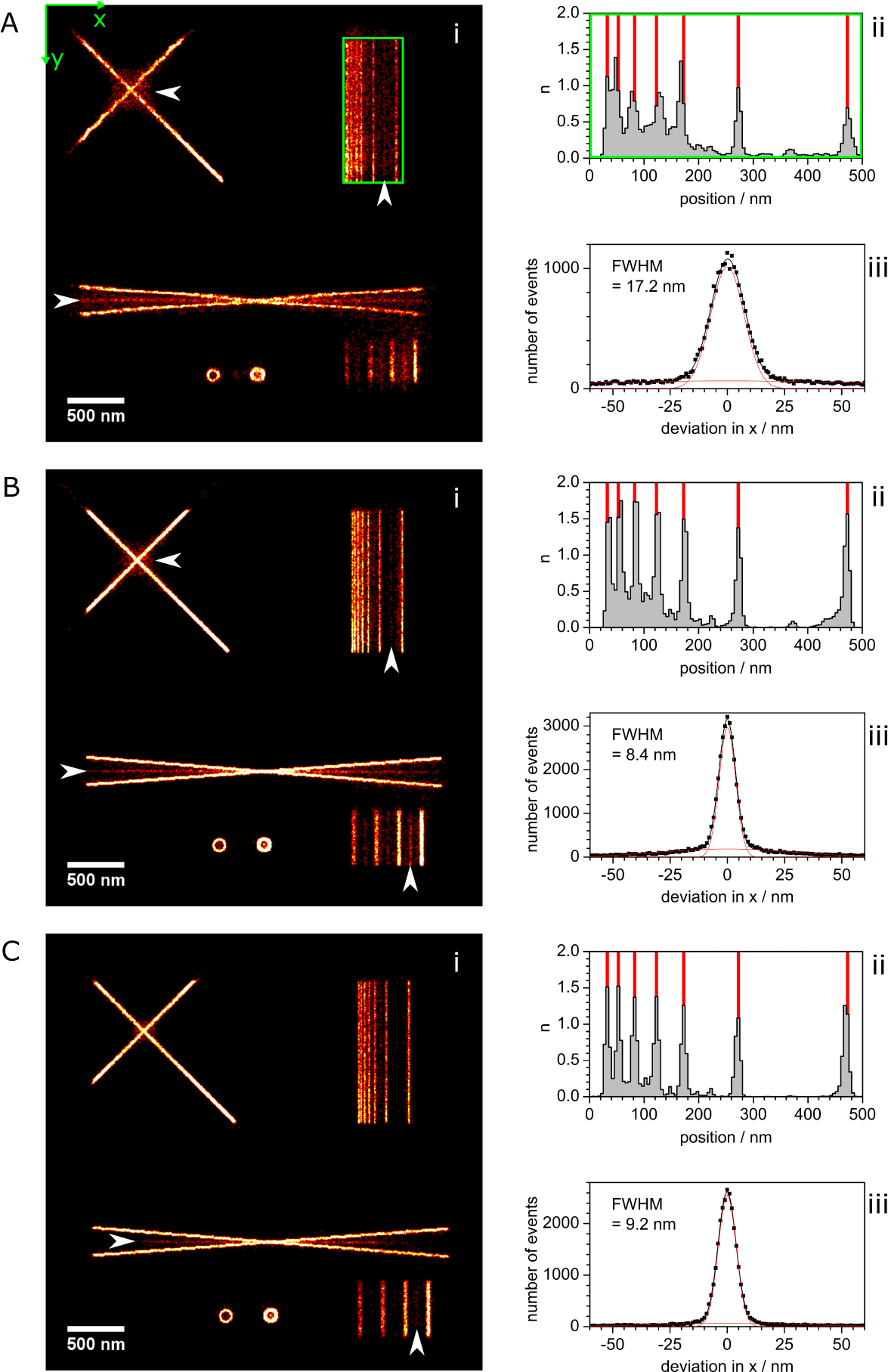
Comparison of the reconstructions by (A) QuickPALM, (B) ThunderSTORM and (C) FIRESTORM. The reconstructions are shown in (i). Apparent image artifacts are marked by white arrows. In (ii) a cross section along the seven parallel line region (indicated by the green square) is shown. True line position are marked in red. Also shown are the histograms displaying the deviation of the localizations from their true position along the x-axis (iii). The distribution can be described by a double Gaussian distribution, shown in black. The single Gaussians are displayed in red.

As mentioned in the statistical analysis, a very high correctness of over 99% was achieved in all three reconstructions. The occurrence of this multitude of reconstruction artifacts is therefore unexpected. However, the observed artifacts occur in the vicinity (*<* 180 nm) of the ground truth structures and are thus not considered a false localization. To quantify the artifacts we determine the deviation between reconstruction and ground truth. Instead of calculating a single value (RMSD), we display the complete histogram of deviations between the ground truth event and the corresponding localization along the *x*-axis (*x*^Test^−*x*^Ref^). As expected, the distribution is centered around 0 which indicates that there is no macroscopic bias. Remarkably, the histogram cannot be fitted by a single gaussian, but two gaussians are needed. One gaussian is rather narrow and has a high amplitude peak and the second gaussian is broad and has a low amplitude. Both are centered around 0 nm deviation as shown in Fig 2. The double Gaussian is thus described by

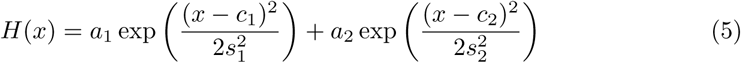

The high amplitude peak is the distribution of correctly localized blinking events. Assuming an appropriate fitting algorithm the width of this peak is ultimately only limited by the Cramér-Rao lower bounds of each individual localization [41]. The second Gaussian has a low amplitude and a broad distribution in comparison. This distribution is a quantification of the observed artifacts in the reconstructions. Based on equation 5 we calculate the ratio Φ of localizations *N*_1_ in the high, narrow Gaussian to the overall number of localization:

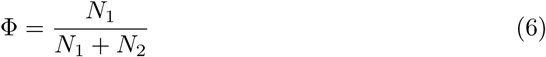

where *N*_2_ is the number of localizations in the low, broad Gaussian. The values *N*_1_, *N*_2_ and Φ for the different reconstructions can be found in table 2. The Φ-value correlates much better with the appearance of artifacts than the RMSD does, proving that the low amplitude, broad Gaussian contains the localizations involved in artifact formation. While ThunderSTORM has 7% more localizations within the narrow gaussian than FIRESTORM (also leading to a reduced FWHM), this comes at the expense of 2.6 times more localizations in the broad gaussian. As a result the Φ-value is reduced by 27% and artifacts appear more prominently in the ThunderSTORM reconstruction as compared to the FIRESTORM reconstruction.

To support the two-population interpretation we plotted the FIRESTORM deviation from the ground truth position in a 2D histogram with the respective number of photons (Fig 3). As described earlier, the blinking events have an expected photon number of 3000 photons. As one would anticipate, we find a narrow main peak at this number of photons which corresponds to the high amplitude Gaussian observed before. We find a second broad localization distribution at around 6000 photons, probably corresponding to the low amplitude Gaussian in Fig 5.

**Table 2.** Number of localizations *N*_1_ in the narrow, high peaked Gaussian population, number of localizations *N*_2_ in the broad, low peaked Gaussian population and the ratio Φ of *N*_1_ to the overall number of localizations for the reconstructions of QuickPALM, ThunderSTORM and FIRESTORM. The underlying simulation contains 99708 blinking events and is done without adding lateral drift.

| | $N_1$ | $N_2$ | $\Phi$ |
| --- | --- | --- | --- |
| QuickPALM | 18451 | 11148 | 0.62 |
| ThunderSTORM | 26681 | 15076 | 0.64 |
| FIRESTORM | 25026 | 5724 | 0.81 |

**Fig 3.**
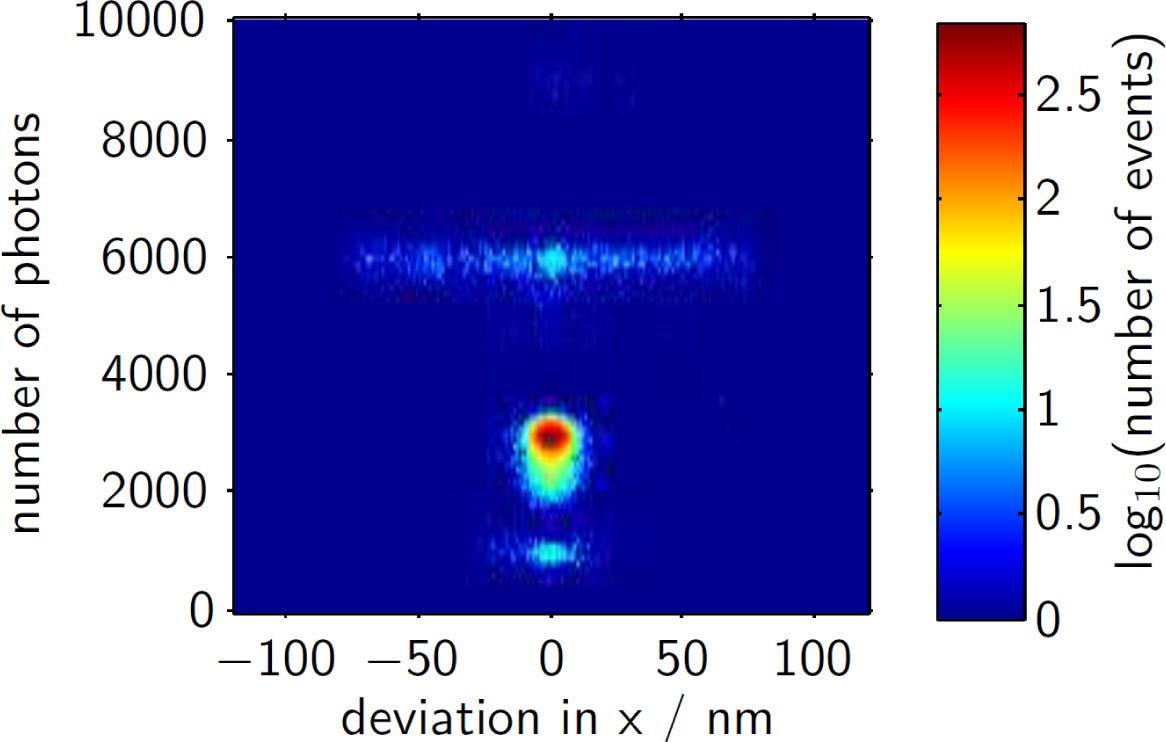
2D Histogram displaying the number of localization events, color coded on a logarithmic scale and plotted over the one dimensional deviation from the ground truth position and the respective number of photons. The main population is centered around the true position and a number of 3000 photons. Additionally one can see a broadly distributed population at 6000 photons associated with artifact formation.

The localizations with a twofold increase in photon count indicate that these artifact-associated localizations originate from the simultaneous emission of two close-by molecules which would also explain the poor average precision of these localizations. This observation offers an explanation for the formation of the ghost lines and dots as well. Even in sparsely labelled samples like the one simulated here, coincidence of multiple blinking events within a diffraction limited area cannot be avoided entirely. In single linear structures emission from sources with overlapping diffraction patterns leads to an apparent localization within the structure. Hence, they do not give rise to apparent artifacts, even though they are mislocalized. Only in regions where the spacing between structures is very short or where structures cross each other the overlap can give rise to blurring effects or ghost structures. For the latter case the intensity is peaked in the center between the ground truth structure (assuming homogeneous labelling density). Therefore, the parameter Φ can be used as metric to determine a SMLM analysis programs ability to avoid reconstruction artifacts caused by the simultaneous localization of overlapping diffraction patterns. Overlapping patterns that are too far apart from each other (i.e. the localization position is farther away from any ground truth position than required by the evaluation threshold of 180 nm) are not counted as matches anymore. Thus, they do not influence the Φ value, instead they lower the value of recall and the Jaccard index.

This kind of artifacts associated with PSF overlap is well known in the SMLM literature [29] [31]. A possible way to remove these artifacts could be to set an upper threshold for the signal photon count. However, in real experiments the signal count does not necessarily scale linearly with the number of emitters even when identical emitters are used. Different axial positions, emitter orientation, background and light scattering in cells or tissue make an adequate choice of an upper threshold infeasible. In particular one does not want to exclude single emitters with a high collected photon number since those can be localized with a very high precision. Another approach is to explicitly incorporate multi-emitter fitting into the localization algorithm which is typically done within the framework of maximum likelyhood estimations or deep learning approaches [42] [43] [44] [22].

### Artifact formation in the presence of drift and inhomogeneous background

Drift is a major problem in single molecule localization which can be reduced using a careful design of the optical setup but not entirely avoided. It usually originates from thermal expansion and contraction of mechanical elements of the setup due to finite temperature fluctuations in the laboratory. Also mechanical vibrations and movements of the sample itself can play a role. Considering the high spatial resolution and the long acquisition time in SMLM, one has to correct for drift during image reconstruction to gain meaningful results. Here, we only simulate lateral drift i.e. drift in *x* and *y*. From a mechanical perspective, there is of course no difference between lateral and axial drift but axial drift can easily be corrected for directly during data acquisition by an appropriate auto-focus system [45]. To illustrate the importance of drift correction we reconstructed the simulated data in presence of non-linear drift (Materials and Methods) and without drift correction as shown in Fig 4. Only the largest spatial structures of the ground truth image (like the large cross in the upper left part of the image) are identifiable while finer structures are lost completely.

**Fig 4.**
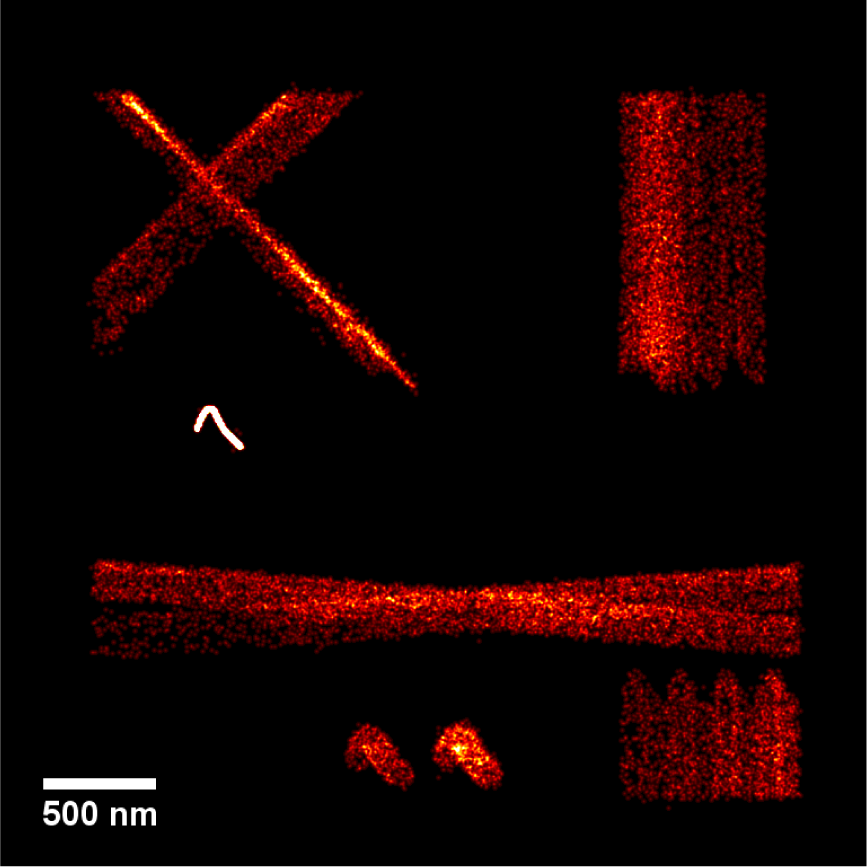
Reconstruction of a simulated data set with strong lateral drift without the application of drift correction. Progression of drift indicated by the course of the fiducial marker left of the center.

The two most common approaches for drift correction are the usage of fiducial markers [35] and (redundant) cross correlation [46] [34]. We compare the reconstructions from ThunderSTORM and FIRESTORM for fiducial marker based and cross correlation based drift correction. QuickPALM performs drift correction only via fiducial markers and is therefore not included in the further analysis. In Fig 5 one can see the drift curves for the different reconstructions, meaning the true lateral position change in black and the drift correction in *x* (red) and *y* (green) estimated by the software tools either by the fiducial marker position or the (redundant) cross correlation.

**Fig 5.**
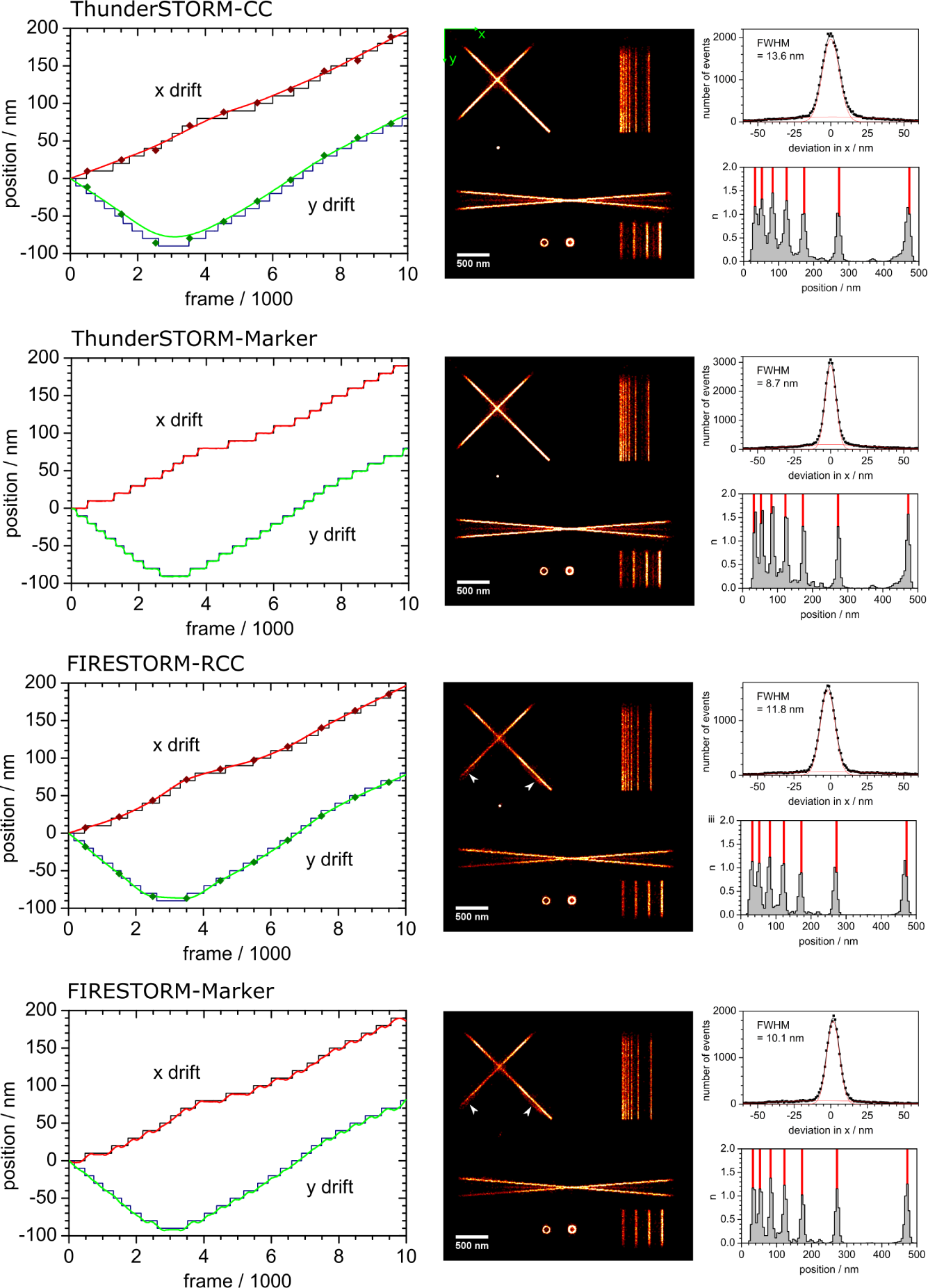
Drift correction curves and reconstruction comparisons of ThunderSTORM and FIRESTORM using (redundant) cross-correlation and fiducial markers. On the left: Drift correction curves in x-direction (red) and y-direction. Ground-truth drift indicated in black. Comparison between ThunderSTORM and FIRESTORM using (redundant) cross correlation as well as fiducial marker. On the right: The corresponding reconstructions, deviation distributions and cross sections. Compare with figure 2

The redundant cross correlation seems to follow the drift more closely than the non-redundant approach. This is in good agreement with earlier observation [34]. Using fiducial markers, the drift correction follows every step very closely, especially in ThunderSTORM. In this distortion free scenario where we simulate drift only as a lateral translation and use a single fiducial marker to infer that translation, the drift correction accuracy only depends on the localization precision with which we identify the true marker position. Thus, we can assume the fiducial marker approach to be superior when it comes to drift correction in this admittedly artificial situation. Unfortunately, adding markers makes experiments more cumbersome. Also, a bright, permanently emitting marker introduces a strong source of inhomogeneous background within the sample. This manifests itself in another type of reconstruction artifacts. In close proximity localizations are biased towards the background peak, meaning the fiducial marker. This is nicely visible in the FIRESTORM reconstructions in the lower arms of the big cross (white arrows in Fig 5).

To correct for this constantly elevated background, one can use a background filter like the temporal running median filter as proposed by Hoogendoorn *et al.* [33]. It uses the fact that blinking events in SMLM typically take place within only a few frames. Thus, they do not change the median intensity of a single pixel over a sufficiently long temporal interval. Subtracting this median intensity hence removes a spatially arbitrary background without interfering with the blinking events themselves as long as that arbitrary background changes sufficiently slow. In Fig 6 the FIRESTORM reconstruction with redundant cross-correlation and temporal running median filter is shown.

**Fig 6.**
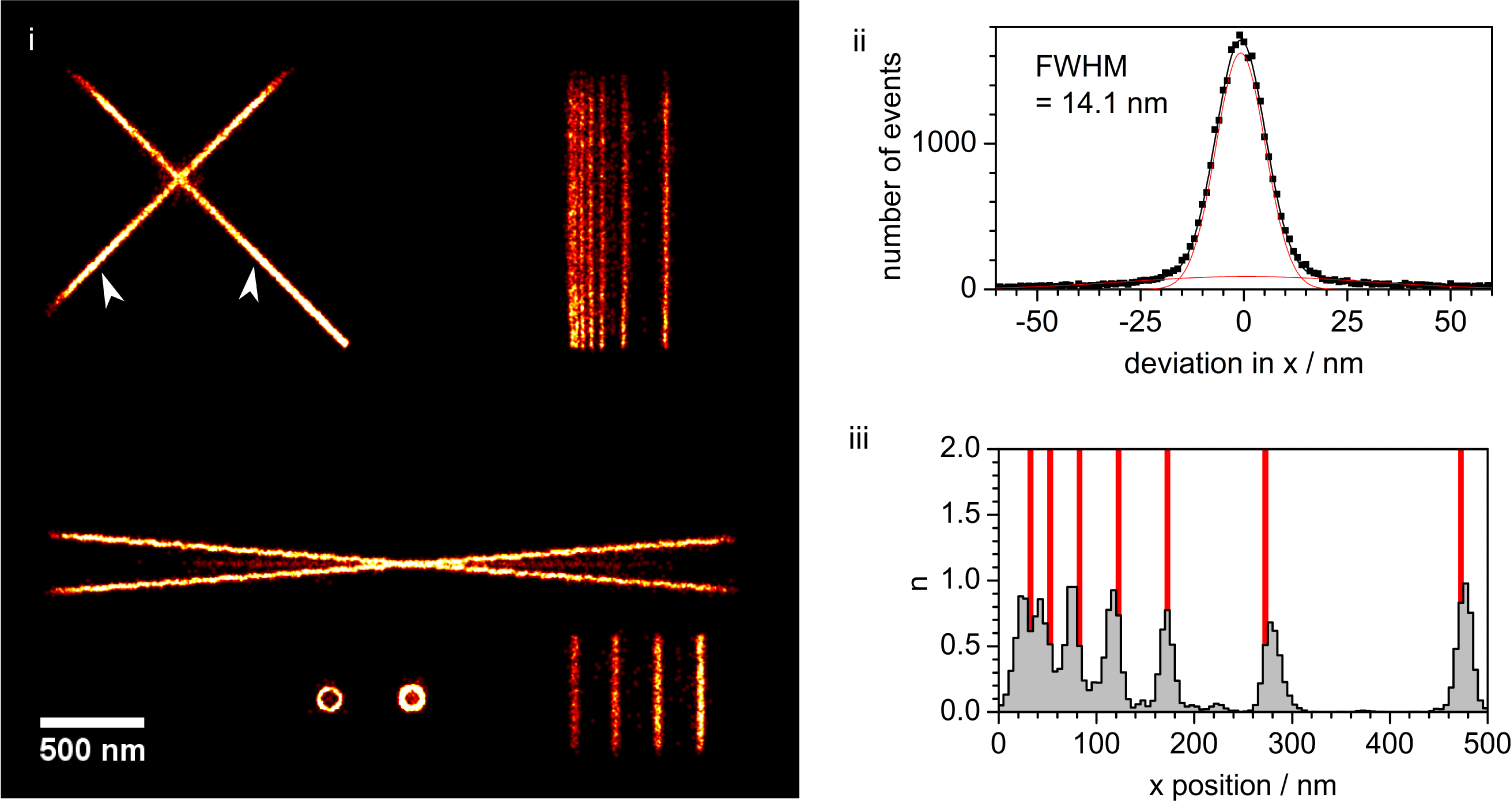
Background (temporal running median filter) and drift (redundant cross-correlation) corrected reconstruction in FIRESTORM (i), Deviation from ground truth positions (ii) and inset showing the region of lines with decreasing spacing distance (iii).

The artifacts induced by the inhomogeneous background due to the fiducial marker are indeed removed as indicated by the white arrows. However, the precision and the trueness of the reconstruction suffer from the background correction. The worse precision is indicated by the increase of the FWHM of the main Gaussian population. The presence of bias is nicely visible in Fig 6 iii: the localizations are biased away from the center of the parallel line structure. Apparently the background correction is particularly problematic in regions with high labelling density. Continuous blinking within this region is misinterpreted by the running median filter as constantly elevated background. This limits the usability of this background correction technique to sparsely labelled samples.

## Conclusion

Artifact formation which is predominantly caused by PSF overlap is a problem in samples of fine spatial structure. Since the ability to resolve structures far beyond the classical resolution limit is the defining characteristic of super resolution in general and SMLM in particular, artifact formation is a serious issue in this field. By comparing the reconstructions of three different SMLM localization and reconstruction tools (QuickPALM, ThunderSTORM and FIRESTORM) we showed that the RMSD value alone is insufficient for the characterization of reconstruction artifact occurrence. Localizations can be grouped into two populations: a group that is truly based on single emitter blinking and a group that is affected by PSF overlap, thus giving rise to several reconstruction artifacts. We introduce a new parameter Φ, the relative fraction of this second population on the entirety of localizations. Φ constitutes an effective measure for the presence of reconstruction artifacts. Thus, we conclude that an appropriate SMLM analysis tool not only needs to perform localization with a high precision, it also has to yield a value of Φ close to unity.

In the absence of drift, PSF overlap is the dominant reason for reconstruction artifact formation. Drift and inhomogeneous background introduce additional artifacts like the bias of localizations along the background intensity gradient. We showed that a widespread background correction algorithm in SMLM indeed restores reconstructed features that were distorted by inhomogeneous background but it might impair the accuracy of reconstructions in densely labelled samples due to the non-negligible time-averaged signal in these areas compared to background.

